# Extended Brain Sources Estimation via Unrolled Optimization Neural Network

**DOI:** 10.1101/2022.04.11.487935

**Authors:** Meng Jiao, Xiaochen Xian, Georges Ghacibeh, Feng Liu

## Abstract

Electroencephalography (EEG)/Magnetoencephalography (MEG) source imaging aims to seek an estimation of underlying activated brain sources to explain the observed EEG/MEG recording. Due to the ill-posed nature of inverse problem, solving EEG/MEG Source Imaging (ESI) requires design of regularization or prior terms to guarantee a unique solution. Traditionally, the design of regularization terms is based on preliminary assumptions on the spatio-temporal structure in the source space. In this paper, we propose a novel paradigm to solve the ESI problem by using Unrolled Optimization Neural Network (UONN) (1) to improve the efficiency compared to traditional iterative algorithms; (2) to establish a data-driven way to model the source solution structure instead of using hand-crafted regularizations; (3) to learn the hyperparameter automatically in a data-driven manner. The proposed framework is based on unfolding of the iterative optimization algorithm with neural network modules. The proposed new learning framework is the first one that use the unrolled optimization neural network to solve the ESI problem. The newly designed framework can effectively learn the source extents pattern and achieved significantly improved performance compared to benchmark algorithms.

## 1 Introduction

Understanding the complex firing neurons and interactions between neural circuits at different brain regions paves an important path to uncover the brain mechanism and brain dysfunctions [1]. Electroencephalography (EEG) or Magnetoencephalography (MEG) is a measurement of the electric potentials on the scalp generated from the electric current sources in the brain [2]. The EEG/MEG measurement is a measures directly the electric firing pattern in the brain, while fMRI, measures the blood-oxygen-level-dependent (BOLD) signal, which is a secondary measurement of the metabolic signal. EEG/MEG has a very high temporal resolution up to 1 millisecond compared to the temporal resolution of around 1 second for fMRI. EEG or MEG devices also have the advantage of being inexpensive, easity portability and versatility. EEG is accepted as a powerful tool to capture the instantaneous brain functionality by measuring the neuronal processes [3]. However, one disadvantage of EEG is its poor spatial resolution and it measures the electric potential on the scalp instead of the underlying sources in the brain. EEG/MEG source imaing (ESI) bridges the gap between the scalp EEG measurement to the brain source activations as it infers the brain sources activation by solving the inverse problem based on the measurement of EEG or MEG [4]. However, given that the dimension of source signal significantly outnumbers the EEG/MEG sensors, the ESI is an ill-conditioned inverse problem that requires sophisticated design of regularizations that utilize the spatial-temporal assumptions on the source space [5, 6].

Recently, numerous algorithms have been developed with different assumptions on the source structure. One seminal work is minimum norm estimate (MNE) where *ℓ*_2_ norm is used as a regularization [7]. Variants of MNE algorithm include dynamic statistical parametric mapping (dSPM) [8] and standardized low-resolution brain electromagnetic tomography (sLORETA) [9]. The *ℓ*_2_-norm based methods tend to render spatially-diffuse source estimation. To promote a sparse solution, Uutela *et al*. [10] introduced the *ℓ*_1_-norm, known as minimum current estimate (MCE). Also, Rao and Kreutz-Delgado proposed an affine scaling method [11] for a sparse ESI solution. Bore *et al*. proposed to use the *ℓ*_*p*_-norm regularization (*p <* 1) on the source signal and the *ℓ*_1_ norm on the data fitting error term [12]. Babadi *et al*. [13] demonstrated that sparse distributed solutions to event-related stimuli can be found using a greedy subspace-pursuit algorithm. It is worthnoting that the sparse constraint can be applied to the orignal source signal or the transformed spatial gradient domain [14, 15]. As the brain is activated not discretely or pointwisely, an extended area of source estimation is prefered [16], and it has been used for multiple applications, such as somatosensory cortical mapping [17], and epileptogenic zone in focal epilepsy patients [18].

Most of the recently developed algorithms requires an iterative procedure to reach the final solution, which can be time consuming. Inspired by the recent advancement of unrolled optimizaiton in solving the inverse problem [19–21], we attempt to use the unrolled optimization deep learning framework to solve the ESI problem to improve the accuracy and effiency solving ESI problem. The advantage of using the unrolled optimization framework is to learn a data-driven regularization instead of using hand-crafted one such as total variation [22], and also replace the iterative procedure with neural network modules, thus improving the online reconstruction efficiency significantly.

## 2 EEG/MEG source imaging problem

EEG data are mostly generated by pyramidal cells in the gray matter with an orientation perpendicular to the cortex. The ESI forward model can be expressed as *Y* = *LS* + *E*, where *Y* ∈ ℝ^*C×T*^ is the EEG/MEG measurements, *C* is the number EEG/MEG channels, *T* is the number of time points, *L* ∈ ℝ^*C×N*^ is the *leadfield* matrix which characterizes the mapping from brain source space to EEG/MEG channel space, *S* ∈ ℝ^*N ×T*^ represents the electrical potentials in *N* source locations for all the *T* time points, and *E* is the uncertainty/noise. The ESI inverse problem is to estimate *S* given the EEG/MEG measurements. Since channel number *C* is much smaller than the number of sources *N*, estimating *S* becomes ill-posed and has infinitely number of solutions. In order to find a unique solution, different regularizations were introduced by using prior assumptions of the source solution. More specifically, *S* can be obtained by solving the following minimization problem:

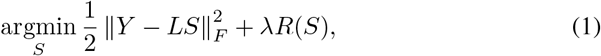

where ∥ · ∥_*F*_ is the Frobenius norm. The first term of Eq.(1) is called *data fitting* which tries to explain the observed EEG data, and the second term is called *regularization* term which is imposed to find a unique solution of Eq.(1) by using the sparsity or other neurophysiology inspired regularizations. If *R*(*S*) equals *ℓ*_2_ norm, the problem is called minimum norm estimate (MNE) [7]; if *R*(*S*) is defined using *ℓ*_1_ norm, the problem becomes minimum current estimate (MCE) [10].

### ESI model with edge sparse total variation

As the cortex is discretized with 3D meshes, simply employing *ℓ*_1_ norm on *S* will result in an estimated descrete source located across the cortex instead of an extended continuous area in the cortex. In order to encourage source extents estimation, Ding proposed to use a sparse constraint in the transformed domain by introducing TV defined from the irregular 3D mesh [22]. Other researchers used the same TV definition such as [4, 23–25]. The TV was defined to be the *ℓ*_1_ norm of the first order spatial gradient using a linear transform matrix *V* ∈ ℝ^*P ×N*^ with its definition can be found in [22], where *N* is the number of voxels/sources, *P* equals the sum of the degrees of all source nodes. Qin et al [26] used a fractional-order total variation term to promote “Gaussian” shape of activation. The model with sparsity and total variation constraints are given as follows:

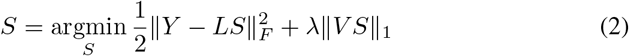

where ∥ · ∥_1_ represents *ℓ*_1_ norm on both row and column of a vector or matrix. The first term is the data fitting term, and the second term is the total variation term. Ideally, the TV regularization promotes source extents estimation. However, it is worthnoting that the TV transormation matrix is *hand-crafted*, either defined on the first or seond spatial derivative, or using fractional-order TV, it can be limiting the flexibility of source configurations. In this paper, we try to *learn* the total variation in a data-driven way by using Unrolled Optimization Neural Network (UONN). The proposed network architecture is given in Fig.1.

**Fig. 1:**
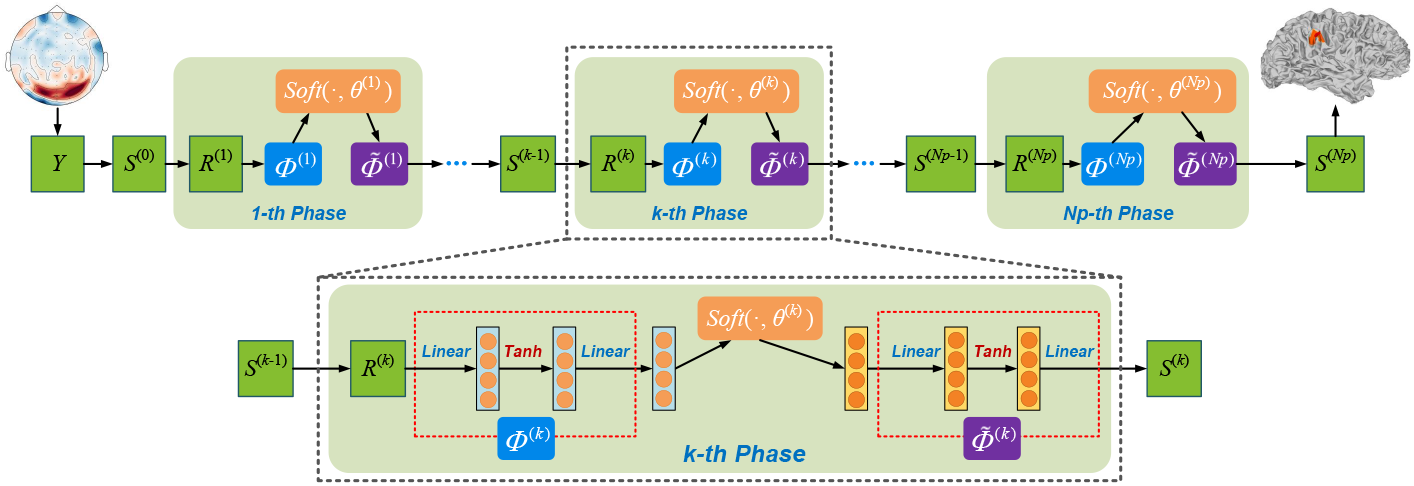
Illustration of the proposed network architecture.

## 3 Proposed unrolled optimization neural network

In this study, we use the recent development of network based compressive sensing (CS) approach [19], the framework is based on iterative shrinkage-thresholding algorithm (ISTA) for solving generally *ℓ*_1_ norm CS problem [27]. The traditional ISTA iterative reconstruction into a deep neural network. To solve Eq. 2, the following update steps are iterated in ISTA:

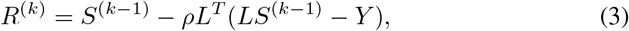

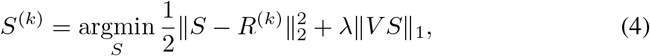

where *k* denotes the interation index, and *ρ* is the step size. Eq. (3) is the gradient decent update of *S*, with an updated *S* denoted as *R*. Eq. (4) is a special case of promximal mapping, *i*.*e*., prox_*λΦ*_(*R*^*K*^) and. Instread of using the hand ∥*V S*∥_1_, we use *Φ*(*S*) to learn the implicit *V S* in a data-driven manner. Thus, the above objective function is re-written as:

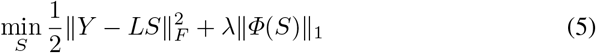

Solving Eq. (5) using ISTA, the Eq. (4) becomes

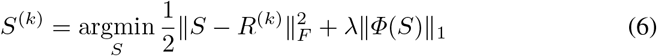

The two iterative steps of Eq.(3) and Eq. (6) can be cast into two seprate modules in the *k*-th iteration of UONN, which is the *R*^(*k*)^ module and *S*^(*k*)^ module. Similar to Zhang & Ghanem [19], we allow the step size *ρ* to vary across iterations, so the output of this module is given as:

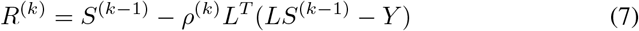

The *S*^(*k*)^ module is to compute the *S*^(*k*)^with an input *R*^(*k*)^. We use an approximated optimization function defined as follows:

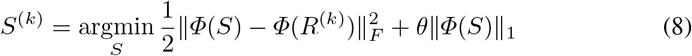

The above equation has a closed-form solution for *Φ*(*S*^(*k*)^), which is

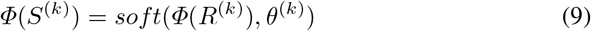

As our goal is to find *S* rather than *Φ*(*S*), we introduce a learnable implicit inverse function 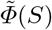, satisfying 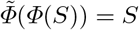. As a result, the update on *S*^(*k*)^ is written as:

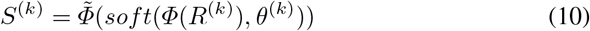

The soft threshold function *soft*(*x, a*) is defined as sign(*x*) max(|*x*| − *a*, 0). The update on *S* is implemented using 2-layers fully connected network connected with a sigmoid activation function (to learn *Φ*(·)), and same struture for 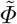, and linked by a soft threshold operator. The structure of proposed network is illustrated in Fig.1.

With this learnable spatial structure expression for *S* using *Φ*(*S*), we aim to learn a more flexible data-driven expression for the extended source activation. The learnable parameters include 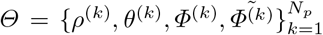, where *N*_*p*_ is the total number of UONN phases, where each phase corresponds to each iteration of ISTA algorithm, and each module in each phase corrsponds to the *R* update and *S* update in Eq. (3) and Eq. (4).

### Loss function

With the training data tuples {*y*_*i*_, *s*_*i*_} for *i* ∈ {1, …, *T* }, the UONN generates the resoncstrution result denoted as 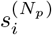, the discrepancy loss is denoted as 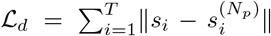, or 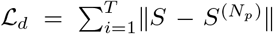. Inspired by the gradual improvement of reconstructed solution, we introduced another smoothness loss 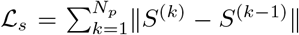, to measure the consistency between two iterative steps to improve the robustness of the algorithm. The final loss function is: ℒ_*d*_ + *α*ℒ_*s*_, and *α* is the weight to balance the above two loss components.

## 4 Numerical Experiments

In this section, we conducted numerical experiments to validate the effectiveness of the proposed method on synthetic EEG data under different Signal Noise Ratios (SNR) settings and also validated it on real data for epileptogenic zone localization. The schematic diagram of the whole process is given in Fig.2.

**Fig. 2:**
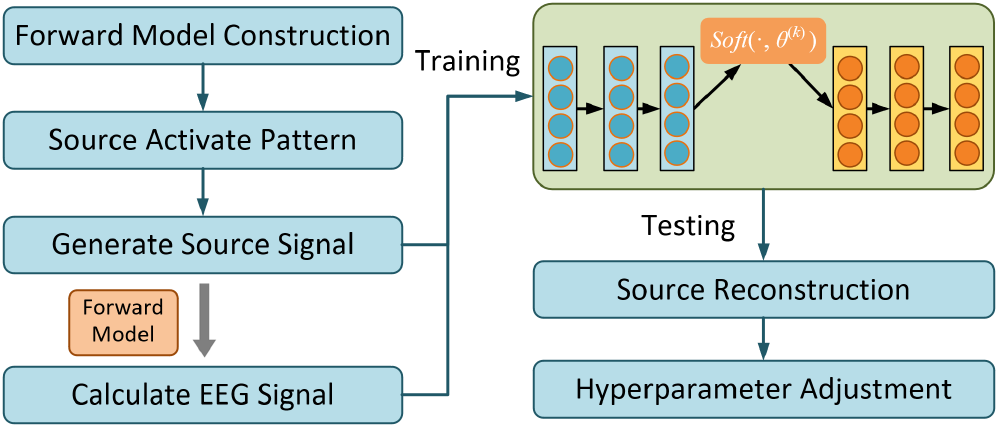
Schematic diagram of the proposed algorithm.

### Simulation experiments

We first conducted experiments on synthetic EEG data with known activation patterns, as the ground truth activation pattern for the

#### Forward model

We used a real head model to calculate the leadfield matrix. The head model was calculated based on T1-MRI images from a 26-year old male subject. Brain tissue segmentation and source surface reconstruction were conducted using FreeSurfer [28]. We used a 128-channel BioSemi EEG cap layout and coregister EEG channels with the head model using Brainstorm and further validated on MNE-Python toolbox [29]. The source space contains 1026 sources in each hemisphere, with 2052 sources combined, resulting in a leadfield matrix *K* with a dimension of 128 by 2052.

#### Experimental settings

To generate EEG data, we activate all 2052 locations in the source space as the central source in turn, and used 3 different neighborhood levels (1-, 2-, and 3-level of neighborhood) to represent different sizes of source extents, illustrated in Fig.3, We activated the whole “patch” of sources corresponding to neighbors at different levels. The activation strength of the central region is set to 1, while the activation strength of the 1-, 2-, and 3-level of adjacent regions is set to 0.8, 0.6, and 0.4 successively.

**Fig. 3:**
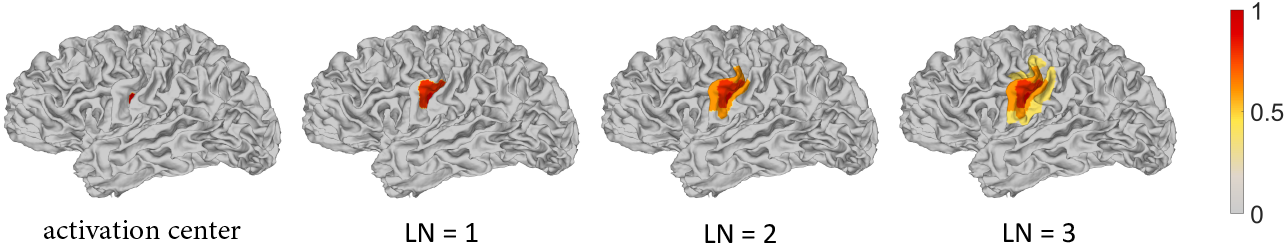
Brain source distributions with different levels of neighbors (LNs).

Then we used the forward model to generate scalp EEG data under different SNR settings (SNR = 40 dB, 30 dB, 20 dB, 10 db and 0 dB). We set the length of EEG data in each experiment setting to be 0.5 second with 100 Hz sampling rate. For each setting of SNR and neighborhood level, we divided the simulated data into training set and testing set according to the proportion of 80% and 20%. The number of layers in the ISTA-net is set to 9, and the *Φ*(·) and 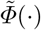 is approximated using a fully connected feed forward neural network with single hidden layer, the input nodes, hidden nodes as well as the output nodes of the network module to approximate *Φ*(·) are set to 2052, 500, and 128, respectively, and the number of input nodes, hidden nodes as well as the output nodes for the network module approximating 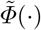 is set to be 125, 500 and 2052. The design of this module archtecture is in resemblace of an autoencoder network [30]. The number of phases is set to be 9.

We chose the training set with 2-level of neighborhood and 40 dB SNR to train the UONN, then the source reconstruction is conducted on all test sets at different levels of source neighborhood configuration and settings of SNR. We picked 10 random source locations to conduct source reconstruction. The benchmark algorithms including MNE [7], dSPM [8], and sLORETA [9] were used for comparison. We quantitatively evaluated the performance of each competing algorithm based on the following metrics:

1. *Localization error (LE)*: it measures the geodesic distance between two source locations on the cortex meshes using the Dijkstra shortest path algorithm.
2. *Area under curve (AUC)*: it is particularly useful to characterize the overlap of an extended source activation pattern.

Better performance for localization is expected if LE is close to 0 and AUC is close to 1. The performance comparison between the proposed method and benchmark algorithms on LE and AUC is summarized in Table 1, and the boxplot figures for SNR=40 dB, 20 dB and 10 dB is given in Fig.4.

**Table 1:** Performance comparison between the proposed method and benchmark algorithms

| SNR | Level of Neighbors (LNs) |  | Source with one LNs |  | Source with two LNs |  | Source with three LNs |  |
| --- | --- | --- | --- | --- | --- | --- | --- | --- |
|  | Method |  | LE | AUC | LE | AUC | LE | AUC |
| 40dB | MNE |  | 37.462 ± 35.318 | 0.938 ± 0.073 | 36.038 ± 34.599 | 0.912 ± 0.076 | 35.813 ± 34.074 | 0.898 ± 0.072 |
|  | sLORETA |  | 12.975 ± 10.221 | 0.963 ± 0.045 | 16.945 ± 11.186 | 0.937 ± 0.041 | 19.426 ± 12.386 | 0.920 ± 0.034 |
|  | dSPM |  | 36.745 ± 26.186 | 0.948 ± 0.045 | 40.136 ± 25.911 | 0.915 ± 0.044 | 42.872 ± 26.083 | 0.892 ± 0.042 |
|  | Proposed |  | **24.937 ± 23.106** | **0.957 ± 0.098** | **6.292 ± 3.155** | **1.000 ± 0.000** | **10.814 ± 6.444** | **0.992 ± 0.017** |
| 30dB | MNE |  | 55.188 ± 54.155 | 0.885 ± 0.116 | 53.664 ± 53.748 | 0.848 ± 0.119 | 55.318 ± 53.923 | 0.826 ± 0.112 |
|  | sLORETA |  | 22.846 ± 30.216 | 0.947 ± 0.060 | 27.358 ± 30.128 | 0.910 ± 0.056 | 29.736 ± 29.882 | 0.884 ± 0.045 |
|  | dSPM |  | 37.027 ± 26.956 | 0.937 ± 0.057 | 40.431 ± 26.631 | 0.894 ± 0.057 | 43.231 ± 26.685 | 0.862 ± 0.056 |
|  | Proposed |  | **24.942 ± 23.066** | **0.957 ± 0.098** | **6.294 ± 3.147** | **1.000 ± 0.000** | **10.834 ± 6.501** | **0.992 ± 0.017** |
| 20dB | MNE |  | 92.003 ± 60.367 | 0.763 ± 0.185 | 99.504 ± 57.910 | 0.716 ± 0.174 | 106.285 ± 54.699 | 0.685 ± 0.158 |
|  | sLORETA |  | 74.682 ± 60.454 | 0.853 ± 0.118 | 82.155 ± 59.295 | 0.791 ± 0.105 | 92.792 ± 57.287 | 0.750 ± 0.091 |
|  | dSPM |  | 40.286 ± 30.014 | 0.869 ± 0.103 | 42.674 ± 29.317 | 0.799 ± 0.095 | 45.388 ± 29.054 | 0.754 ± 0.089 |
|  | Proposed |  | **25.061 ± 23.241** | **0.956 ± 0.100** | **6.410 ± 3.257** | **1.000 ± 0.000** | **10.851 ± 6.618** | **0.992 ± 0.017** |
| 10dB | MNE |  | 115.073 ± 49.547 | 0.593 ± 0.224 | 115.424 ± 48.802 | 0.567 ± 0.196 | 116.015 ± 48.633 | 0.550 ± 0.177 |
|  | sLORETA |  | 111.976 ± 49.858 | 0.631 ± 0.187 | 112.728 ± 48.738 | 0.593 ± 0.151 | 113.480 ± 48.163 | 0.570 ± 0.131 |
|  | dSPM |  | 72.674 ± 49.649 | 0.650 ± 0.172 | 74.898 ± 49.472 | 0.606 ± 0.132 | 77.547 ± 48.895 | 0.580 ± 0.112 |
|  | Proposed |  | **25.861 ± 23.873** | **0.953 ± 0.104** | **7.434 ± 4.059** | **1.000 ± 0.001** | **11.492 ± 7.309** | **0.989 ± 0.021** |
| 0 dB | MNE |  | 116.648 ± 47.874 | 0.515 ± 0.219 | 116.610 ± 47.823 | 0.510 ± 0.195 | 116.950 ± 47.832 | 0.507 ± 0.179 |
|  | sLORETA |  | 114.490 ± 47.332 | 0.520 ± 0.185 | 114.123 ± 47.195 | 0.513 ± 0.155 | 114.392 ± 47.165 | 0.510 ± 0.138 |
|  | dSPM |  | 108.192 ± 48.302 | 0.523 ± 0.169 | 108.843 ± 48.346 | 0.515 ± 0.132 | 107.938 ± 47.561 | 0.511 ± 0.113 |
|  | Proposed |  | **32.706 ± 29.557** | **0.920 ± 0.145** | **15.298 ± 14.230** | **0.986 ± 0.041** | **19.266 ± 17.243** | **0.948 ± 0.065** |

**Fig. 4:**
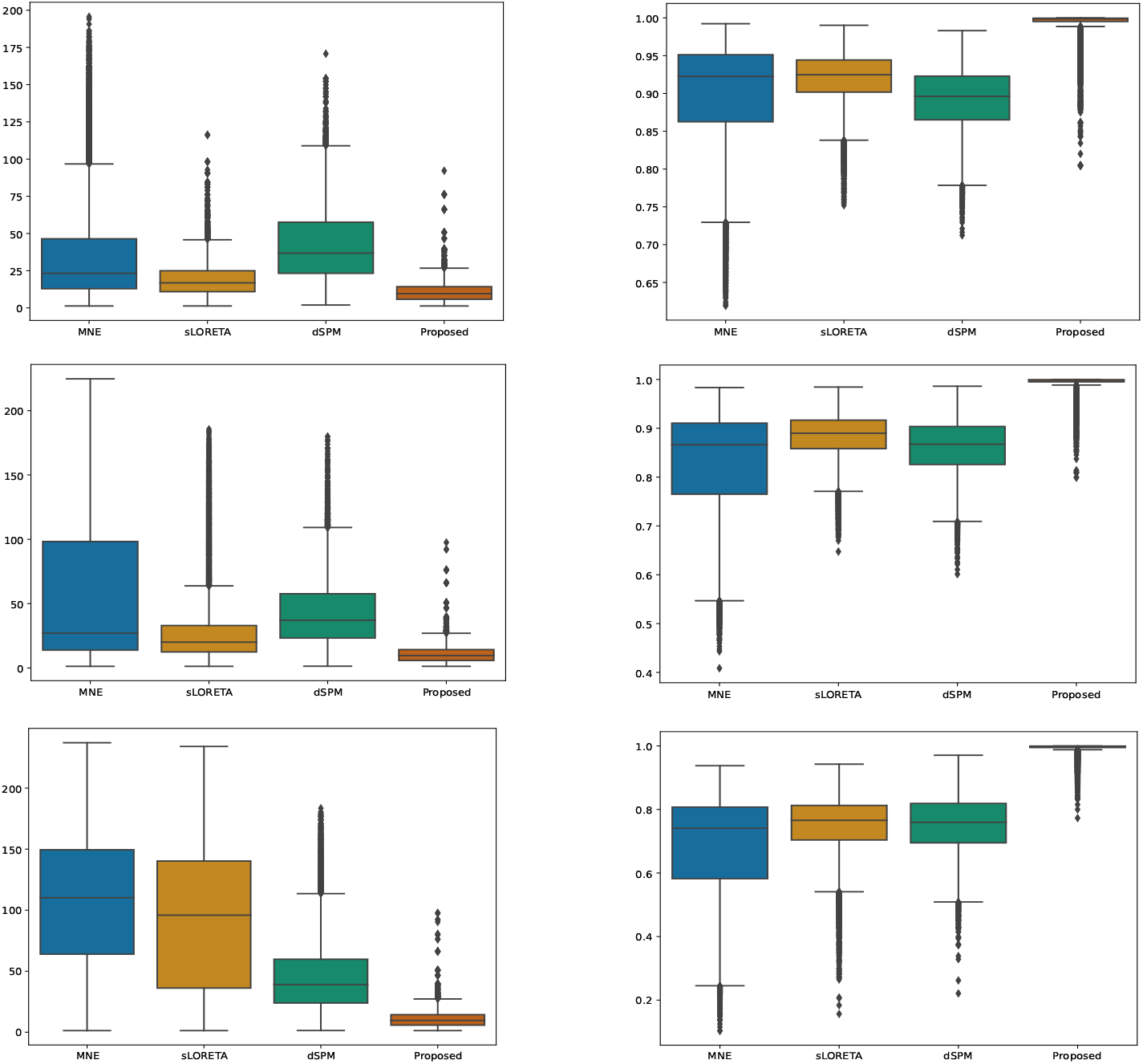
Peformance comparison of different algorithms on LE (on the left side) and AUC (on the right side), for SNR = 40 dB (on the top row), SNR = 30 dB (on the second row) and SNR =20 dB (on the bottom row).

From Table 1, we can see that compared to the benchmark algorithms MNE, dSPM, and sLORETA, the proposed UONN method exhibits excellent performance, while the benchmark algorithms can only provide satisfactory source reconstruction when the activated area is small and the SNR level is low. With the expansion of the source range and the increase of noise level, the LE values of the benchmark algorithms deteriorate significantly, so as to the AUC values. By contrast, the superiority of the UONN method is demonstrated by improved AUC values and smaller LE values. It shows better performance for larger source extents and higher noise level compared to the benchmark algorithms. The corresponding LE values are always at a low level, and the AUC values are always stable above 0.94, which demonstrates the excellent stability of the proposed method. By comparing different source activation size, we can see larger source extents are more difficult to be localized exactly, with a worse AUC then smaller areas of activation, however, the localization error can be lower in the benchmark algorithms. The noise can impact significantly on the performance of all the benchmark algorithms, however, when SNR equal or above 10 dB, our algorithm performs very well in recovering the source extents. The comparison between the reconstructed source distributions based with 3-level of neighborhood and 40 dB, 30 dB and 20 dB SNR is shown in Fig. 5.

**Fig. 5:**
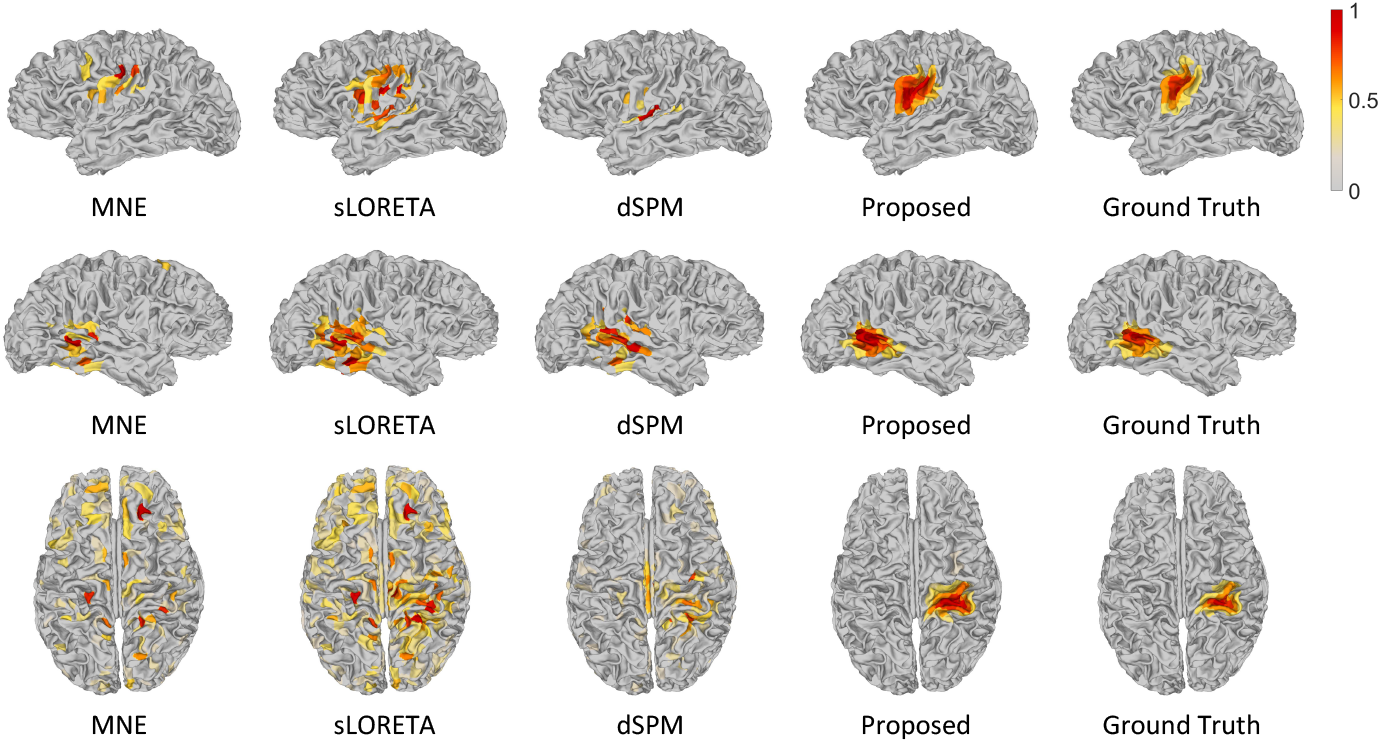
Brain sources reconstruction by different ESI algorithms with 3-level of neighborhood for SNR = 40 dB (on the top row), SNR = 30 dB (on the second row) and SNR =20 dB (on the bottom row).

The reconstructed source distributions based on outputs of different layers is shown in Fig. 6. From this figure, we can see that: Due to the introduction of the smoothness loss ℒ_*s*_, the results obtained at each iteration show significant smoothness. Besides, as the iteration proceeds (the layer index increases), the reconstructed source distribution gradually evolves towards the ground truth.

**Fig. 6:**
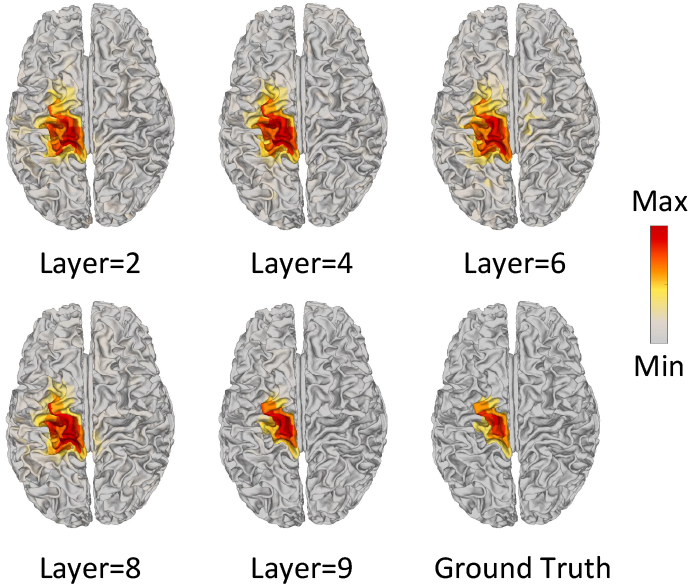
Source distributions based on outputs of different layers.

### 4.1 Real data experiments

We analyzed a real dataset that is publicly accessible through the Brainstorm tutorial datasets [31]. This dataset was recorded from a patient with focal epilepsy who was seizure-free during a 5-year follow-up period after a left frontal tailored resection. We followed the Brainstorm tutorial to obtain the head model and the lead field matrix, then we calculated the average spikes (as shown in Fig. 7) of the EEG measurements of 29 channels. We used the peak (0 ms) of the averaged interictal spike for brain source localization, and the comparison between the reconstructed source distributions based on different methods is shown in Fig. 8.

**Fig. 7:**
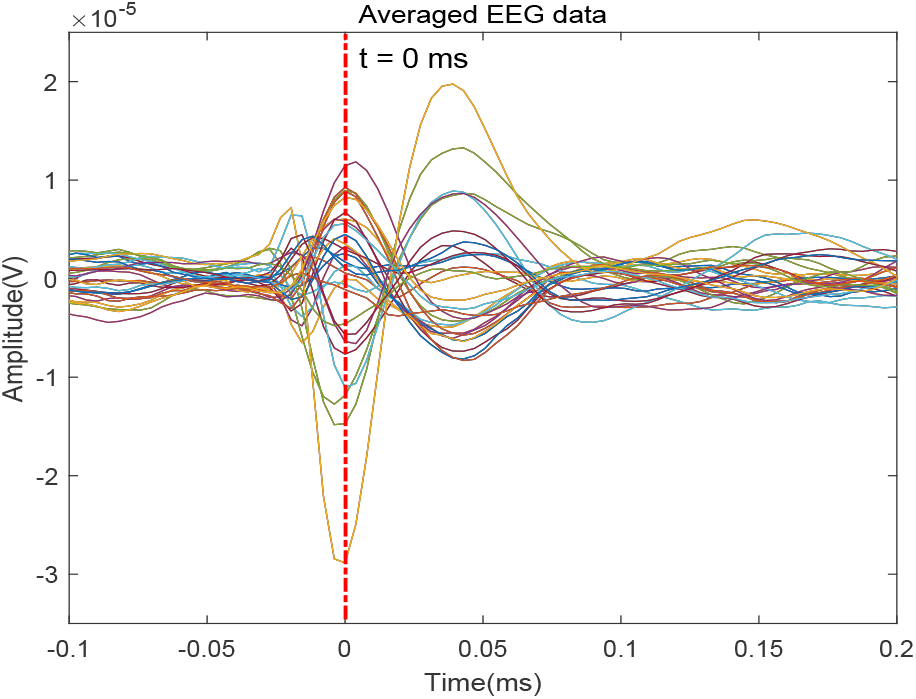
Average EEG time series plot around the inter-ictal spike.

**Fig. 8:**
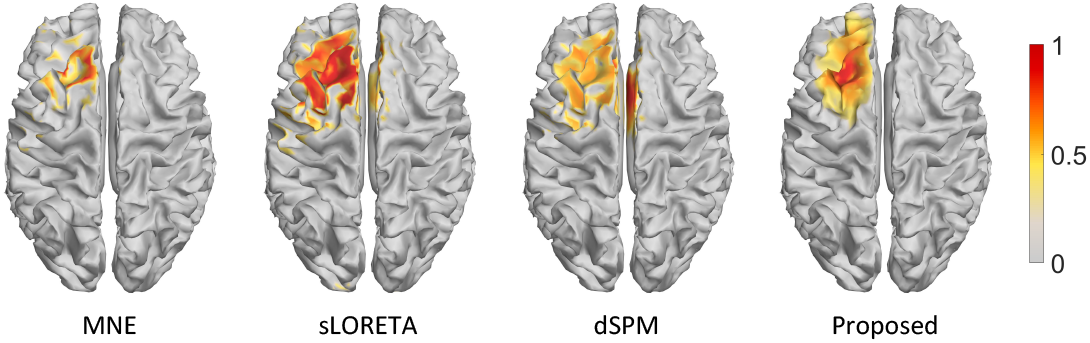
Reconstructed sources by different ESI algorithms for epilepsy EEG data.

From Fig. 8, we can see that both the MNE method and the proposed UONN method can accurately locate the epileptogenic zone, which was validated by the follow-up survey after resection on the left frontal region. From the source reconstruction results of the proposed method, it can be seen that in addition to the accurate location of the lesion area, the difference in signal strength between the central region and its adjacent regions can also be seen clearly. This shows that the UONN method can reconstruct source signals of different intensities corresponding to neighbors of different levels. In contrast, the source distribution area estimated by sLORETA and dSPM is highly broad, although the left frontal region is covered, part of the right frontal which is not related to the epilepsy lesion is also included. Obviously, among these methods, the proposed UONN method provides a cleaner and accurate estimation of the epileptogenic zone.

## 5 Conclusion

In this paper, we propose a new method based on deep learning unrolled optimization framework for brain source reconstruction. The proposed framework enjoys the advantage of great approximation capability and principled parameter training procedure in deep learning. It eliminate the necessity of hand-crafted total variation in the traditional method. We designed a neural network module to learn the spatial structure of source extents. We incorporated a smoothness constraint in the loss function to mimic the iterative changes in the solution. The numerical experiments demonstrated a great boost in the performance measured by AUC and LE.

## Notes

### Competing Interest Statement

The authors have declared no competing interest.

### Summary of Updates

Add more experiments and detailed explanation on the methods.

